# Structural and biochemical characterization of nsp12-nsp7-nsp8 core polymerase complex from COVID-19 virus

**DOI:** 10.1101/2020.04.23.057265

**Authors:** Qi Peng, Ruchao Peng, Bin Yuan, Jingru Zhao, Min Wang, Xixi Wang, Qian Wang, Yan Sun, Zheng Fan, Jianxun Qi, George F. Gao, Yi Shi

## Abstract

The ongoing global pandemic of coronavirus disease 2019 (COVID-19) has caused huge number of human deaths. Currently, there are no specific drugs or vaccines available for this virus. The viral polymerase is a promising antiviral target. However, the structure of COVID-19 virus polymerase is yet unknown. Here, we describe the near-atomic resolution structure of its core polymerase complex, consisting of nsp12 catalytic subunit and nsp7-nsp8 cofactors. This structure highly resembles the counterpart of SARS-CoV with conserved motifs for all viral RNA-dependent RNA polymerases, and suggests the mechanism for activation by cofactors. Biochemical studies revealed reduced activity of the core polymerase complex and lower thermostability of individual subunits of COVID-19 virus as compared to that of SARS-CoV. These findings provide important insights into RNA synthesis by coronavirus polymerase and indicate a well adaptation of COVID-19 virus towards humans with relatively lower body temperatures than the natural bat hosts.

## Introduction

In the end of 2019, a novel coronavirus (2019-nCoV) caused an outbreak of pulmonary disease in China (Zhu et al., 2020), which was later officially named “severe acute respiratory syndrome virus 2” (SARS-CoV-2) by the International Committee on Taxonomy of Viruses (ICTV) (Coronaviridae Study Group of the International Committee on Taxonomy of, 2020). The pneumonia disease was named coronavirus disease 2019 (COVID-19) by world health organization (WHO). The outbreak has developed into a global pandemic affecting most countries all over the world (Holshue et al., 2020; Kim et al., 2020). As of April 19^th^, 2020, more than 2,000,000 human infections have been reported worldwide, including over 160,000 deaths (https://www.who.int/emergencies/diseases/novel-coronavirus-2019). The origin of this virus has not been identified, but multiple origins possibly exist based on the recent bioinformatics analysis of the viral isolates from different countries (Andersen et al., 2020; Zhang and Holmes, 2020). So far, there are no specific drugs or vaccines available yet, which poses a great challenge for the treatment and control of the diseases.

SARS-CoV-2 belongs to the family of *Coronaviridae*, a group of positive-sense RNA viruses with a broad host-spectrum (Vicenzi et al., 2004). Currently, a total of seven human-infecting coronaviruses have been identified, among which SARS-CoV-2 displays the highest similarity in genome sequence to the SARS-CoV emerged in 2002-2003 (Zhong et al., 2003; Zhou et al., 2020). Both viruses utilize the same host receptor angiotensin-converting enzyme 2 (ACE2) for cell entry and cause respiratory symptoms that may progress to severe pneumonia and lead to death (Lu et al., 2020; Zhou et al., 2020). However, as compared to the SARS-CoV, SARS-CoV-2 shows a much higher transmission rate and a lower mortality (Huang et al., 2020; Wang et al., 2020). Most of the infections result in mild symptoms and a substantial number of asymptomatic infection cases have also been reported (Rothe et al., 2020). These properties allow the SARS-CoV-2 to transmit among humans furtively, facilitating the quantum-leaps of pandemic expansion. Characterizing the infection and replication behaviors of SARS-CoV-2 would provide critical information for understanding its unique pathogenesis and host-adaption properties.

The replication of coronavirus is operated by a set of non-structural proteins (nsps) encoded by the open-reading frame 1a (ORF1a) and ORF1ab in its genome, which are initially translated as polyproteins followed by proteolysis cleavage for maturation (Ziebuhr, 2005). These proteins assemble into a multi-subunit polymerase complex to mediate the transcription and replication of viral genome. Among them, nsp12 is the catalytic subunit with RNA-dependent RNA polymerase (RdRp) activity (Ahn et al., 2012). The nsp12 itself is capable of conducting polymerase reaction with extremely low efficiency, whereas the presence of nsp7 and nsp8 cofactors remarkably stimulates its polymerase activity (Subissi et al., 2014). The nsp12-nsp7-nsp8 subcomplex is thus defined as the minimal core components for mediating coronavirus RNA synthesis. To achieve the complete transcription and replication of viral genome, several other nsp subunits are required to assemble into a holoenzyme complex, including the nsp10, nsp13, nsp14 and nsp16, for which the precise functions for RNA synthesis have not been well understood (Adedeji et al., 2012; Lehmann et al., 2015; Sevajol et al., 2014; Ziebuhr, 2005). The viral polymerase has shown enormous prospect as a highly potent antiviral drug target due to its higher evolutionary stability, compared to the surface proteins that are more prone to drift as a result of the selection by host immunity (Shi et al., 2013). Therefore, understanding the structure and functions of SARS-CoV-2 polymerase complex is an essential prerequisite for developing novel therapeutic agents.

In this work, we determined the near-atomic resolution structure of SARS-CoV-2 nsp12-nsp7-nsp8 core polymerase complex by cryo-electron microscopy (cryo-EM) reconstruction, and revealed the reduced polymerase activity and thermostability as compared to its SARS-CoV counterpart. These findings improve our understanding of coronavirus replication and evolution which might contribute to the fitness of SARS-CoV-2 to human hosts.

## Results

### Overall structure of SARS-CoV-2 core polymerase complex

The SARS-CoV-2 nsp12 polymerase and nsp7-nsp8 cofactors were expressed using the baculovirus and *Escherichia coli* (*E. coli*) expression systems, respectively. The three protein subunits were mixed *in vitro* to constitute the core polymerase complex (Figure S1). The structure of SARS-CoV-2 nsp12-nsp7-nsp8 complex was determined at 3.7 Å resolution, which clearly resolved the main chain trace and most bulky side chains of each subunit (Figures S2 and S3). Similar to the counterpart complex of SARS-CoV (Kirchdoerfer and Ward, 2019), the N-terminal ~110 amino acids of nsp12, as well as the N-terminal ~80 residues of nsp8 and a small portion of nsp7 C-terminus, could not be resolved in the density map (Figure 1A). In total, approximately 80% of the 160-kDa complex was interpreted in the structure.

**Figure 1.**
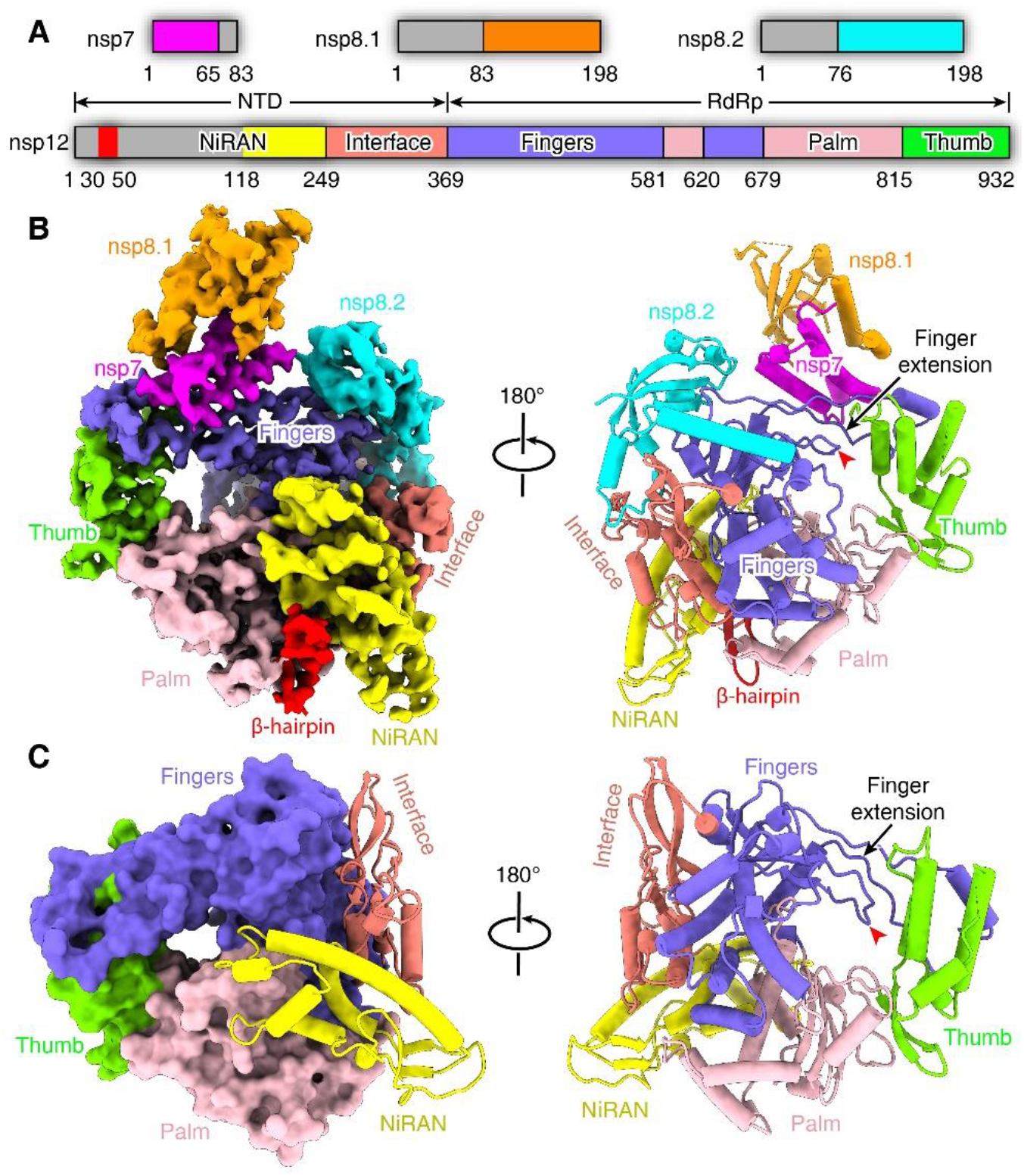
Overall structure of SARS-CoV-2 core polymerase complex. **(A)** Schematic diagram of domain architecture for each subunit of polymerase complex. Each domain is represented by a unique color. The unresolved region is colored in grey. **(B)** Overall density map (left) and atomic model (right) of the SARS-CoV-2 nsp12-nsp7-nsp8 core complex at different views. Both the map and structural model are colored by domains with the same color code as in **(A)**. The finger-tip loop (one of the key catalytic motifs) is highlighted with a red arrowhead, and the associated finger extension loops are indicated by a black arrow. **(C)** The structure of nsp12 polymerase subunit in different views, colored by domains with the same scheme as in **(A)**. **See also Figures S1-S3 and Table S1**.

The SARS-CoV-2 polymerase complex consists of a nsp12 core catalytic subunit bound with a nsp7-nsp8 heterodimer and an additional nsp8 subunit at a different binding site (Figure 1B). The N-terminal portion of nsp12 polymerase subunit contains a Nidovirus RdRp-associated nucleotidyltransferase (NiRAN) domain that is shared by all members of the *Nidovirales* order (Lehmann et al., 2015). This domain binds at the back side of the right-hand configured C-terminal RdRp. Between them is an interface domain that links the NiRAN domain to the fingers subdomain of RdRp (Figure 1B and C). The NiRAN and interface domains represent additional features of coronavirus RdRp as compared to the polymerase subunit of flaviviruses which is also a group of positive-sense RNA viruses (Duan et al., 2017; Godoy et al., 2017; Zhao et al., 2017). The C-terminal catalytic domain adopts a conserved architecture of all viral RdRps, composed of the fingers, palm and thumb subdomains (Figure 1C). A remarkable feature of the coronavirus RdRp is the long finger extension that intersects with the thumb subdomain to form a closed-ring structure (Figure 1C), in contrast to the smaller loop in segmented negative-sense RNA virus (sNSV) polymerases which results in a relatively open conformation, such as influenza virus, bunyavirus and arenavirus polymerases (Figure S4) (Gerlach et al., 2015; Peng et al., 2020; Pflug et al., 2014). Similar close contact between the fingers and thumb subdomains is also observed in the structures of poliovirus (PV) and Zika virus (ZIKV) polymerases (Figure S4) (Godoy et al., 2017; Gong and Peersen, 2010), which might be a common feature of positive-sense RNA virus polymerases.

### Structural comparison of SARS-CoV-2 and SARS-CoV nsp12-nsp7-nsp8 complexes

Basically, the structure of SARS-CoV-2 polymerase complex highly resembles that of SARS-CoV, with a global root mean square deviation (RMSD) of ~1 Å for the α-carbon atoms (Figure 2A). There are 1, 4 and 25 residue substitutions between the two viruses in the structurally visualized regions of nsp7, nsp8 and nsp12 subunits, respectively (Figure 2B) (there are 1 and 7 additional site mutations in the unresolved regions of nsp8 and nsp12 subunits, respectively). However, these mutations did not result in obvious structural changes of the polymerase complex. During the review process of this manuscript, two other research groups also reported the high-resolution structure of SARS-CoV-2 core polymerase complex at atomic resolutions, which revealed similar structural features to the SARS-CoV counterpart, consistent with our observations (Gao et al., 2020; Yin et al., 2020). All of the three structures reported by our group and others’ revealed similar structural features, including the partially unresolved N-terminal portion of nsp8 and a small C-terminal tail of nsp7 subunits, and allowed the identification of a previously undefined β-hairpin motif in the N-terminus of nsp12 subunit which binds at the interface between the NiRAN domain and the palm subdomain of RdRp (Figure 2C). However, no extra density for this region was observed in the reconstruction of SARS-CoV polymerase complex (Figure 2D), suggesting the different conformation or flexibility of this motif between the two viruses. Of note, in the structure determined by Gao et al., the N-terminal residues 51-117 of nsp12 subunit were clearly resolved to constitute an almost complete NiRAN domain (Gao et al., 2020). In contrast, this region revealed poor density in the reconstructions by our group and Yin et al., suggesting the moderate flexibility of this region (Yin et al., 2020), for which the functional relevance yet remained elusive.

**Figure 2.**
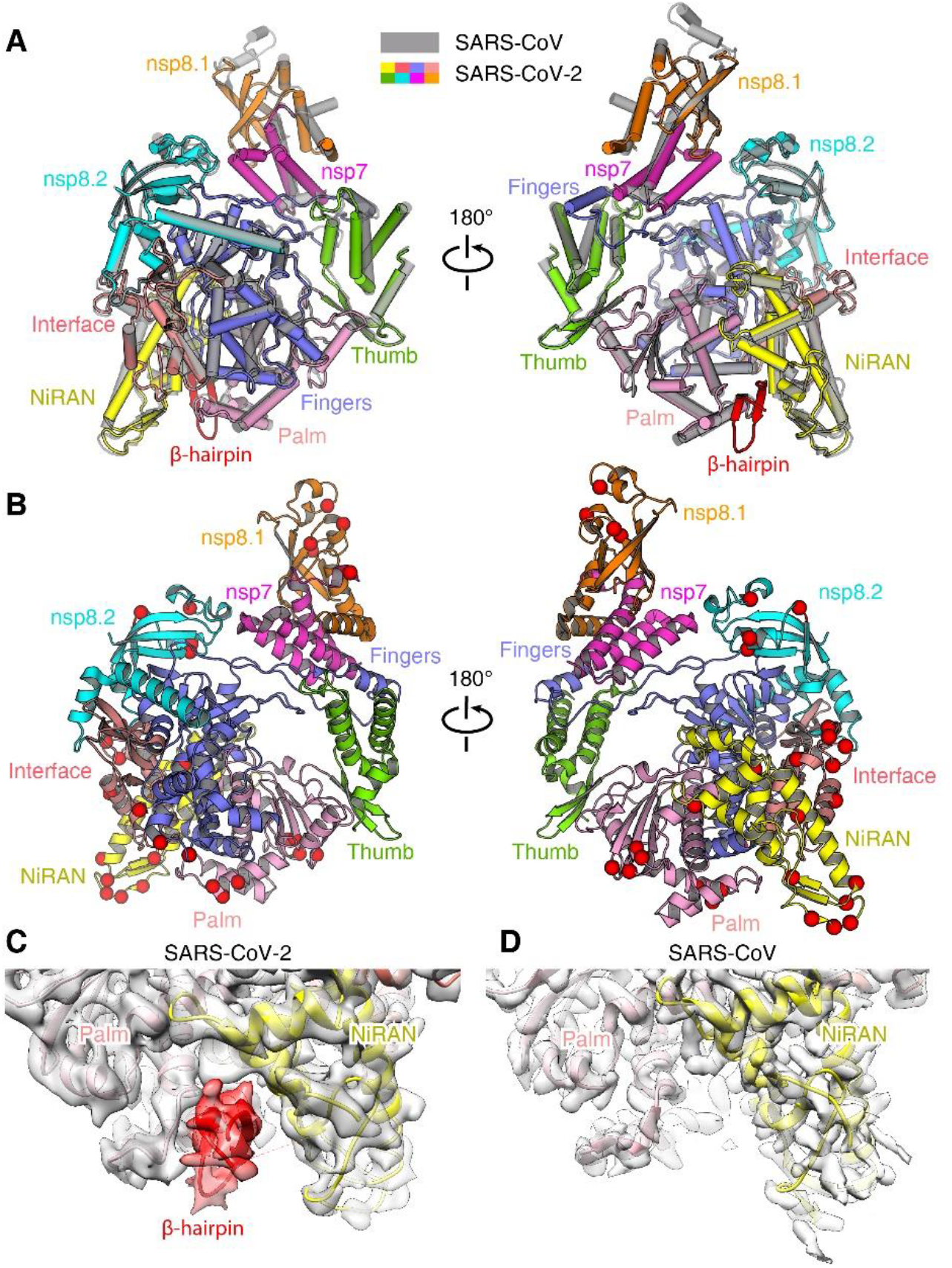
Structural comparison of SARS-CoV-2 and SARS-CoV core polymerase complexes. **(A)** Overlay of the nsp12-nsp7-nsp8 complexes of SARS-CoV-2 (colored by domains) and SARS-CoV (grey). The two structures could be well superimposed with high similarity. **(B)** Residue substitutions between SARS-CoV-2 and SARS-CoV core polymerase complexes. The structural model is shown in cartoons and colored by domains. The substitution sites are represented by red spheres to highlight their locations. **(C-D)** Comparison of nsp12 N-terminal densities in SARS-CoV-2 **(C)** and SARS-CoV **(D)** polymerase complexes. The density for the newly identified N-terminal β-hairpin of SARS-CoV-2 nsp12 subunit is highlighted in red. No corresponding density was observed in the reconstruction for SARS-CoV polymerase complex. **See also Figure S4**.

### Conserved catalytic center of nsp12 and interaction with cofactors

The catalytic domain of SARS-CoV-2 nsp12 subunit is arranged following the typical right-hand configuration shared by all viral RdRps, which includes seven critical catalytic motifs (A-G) (Figure 3). Among them, motifs A-F are highly conserved for all viral RdRps, and the motif G is defined as a hallmark of primer-dependent RdRp in some positive-sense RNA viruses which interacts the primer strand to initiate RNA synthesis (Figure S4). The motif C contains the critical 759-SDD-761 catalytic residues which reside in a β-turn loop connecting two adjacent strands. The motif F forms a finger-tip that protrudes into the catalytic chamber and interacts with the finger extension loops and the thumb subdomain (Figure 3B). It has been shown that some sNSV polymerases, e.g. influenza virus and bunyavirus polymerases, require the binding of a conserved 5’-RNA hook to activate the activity for RNA synthesis by stabilizing the finger-tip which is otherwise highly flexible in the apo form (Gerlach et al., 2015; Hengrung et al., 2015; Peng et al., 2020; Pflug et al., 2014; Reich et al., 2014). In the structure of coronavirus polymerase, this finger-tip loop is stabilized by the adjacent finger extension loops which are secured by the interactions with nsp7-nsp8 heterodimer (Figure 1B and C). In the absence of this heterodimer, the finger extension loops displayed significant flexibility as observed in the structure of SARS-CoV nsp12-nsp8 subcomplex, which would thus destabilize the finger-tip motif (Figure S4G and H) (Kirchdoerfer and Ward, 2019). These evidences are consistent with the observation that the nsp12 alone shows extremely weak activity for nucleotide polymerization, whereas this activity is remarkably stimulated upon the binding of nsp7-nsp8 cofactors (Ahn et al., 2012; Subissi et al., 2014).

**Figure 3.**
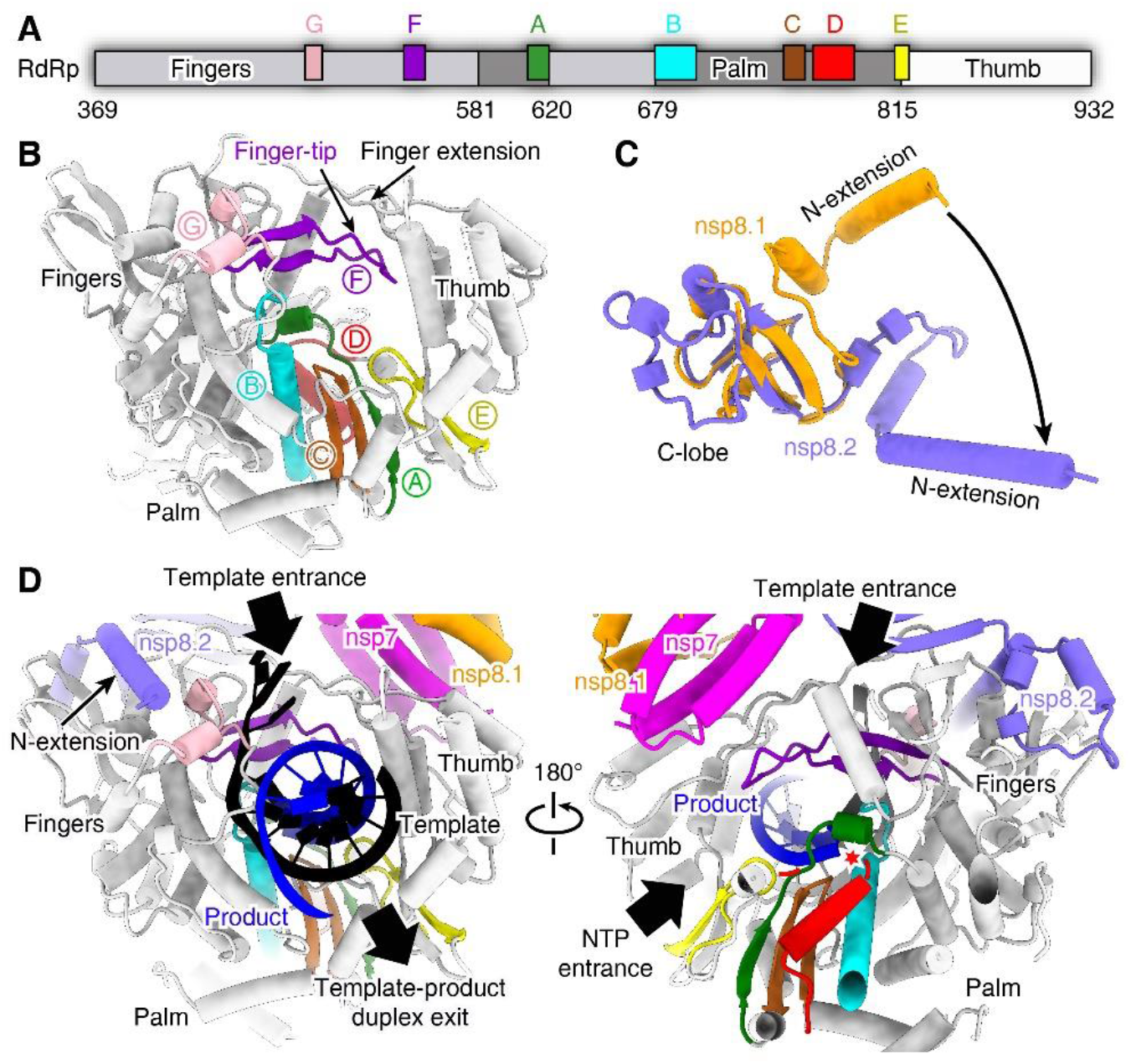
Catalytic center of nsp12 RdRp and the interactions with nsp7-nsp8 cofactors. **(A)** Schematic diagram of domain organization of SARS-CoV-2 nsp12 RdRp region. The RdRp domain consists of fingers, palm and thumb subdomains. The seven conserved catalytic motifs are indicated by different colors in corresponding locations. **(B)** Atomic structure of nsp12 shown in cartoons with the seven catalytic motifs colored differently as in **(A)**. **(C)** Structural comparison of the two nsp8 subunits in the complex. The C-terminal lobe could be well superimposed, while the N-terminal extension helix shows different conformations. **(D)** Structural model for RNA synthesis by coronavirus polymerase complex. The template (black) and product (blue) strands are modeled based on the elongation complex of poliovirus polymerase (PDB ID: 3OL8). The template and NTP entrance, and the template-product duplex exit tunnels are indicated by black arrows. The conserved RdRp catalytic motifs are colored with the same scheme as in **(A)**. The catalytic site is highlighted with a red star. **See also Figures S4 and S5**.

The nsp7-nsp8 heterodimer binds above the thumb subdomain of RdRp and sandwiches the finger extension loops in between to stabilize its conformation (Figure 1B). This interaction is mainly mediated by the nsp7 within the heterodimer while the nsp8 (nsp8.1) contributes few contacts with the nsp12 polymerase subunit (Figures 1B and 3D). The other nsp8 (nsp8.2) subunit clamps the top region of the finger subdomain and forms additional interactions with the interface domain (Figures 1B and 3D). The two nsp8 subunits display significantly different conformations with substantial refolding of the N-terminal extension helix region, which mutually preclude the binding at the other molecular context (Figure 3C). The importance of both cofactor-binding sites has been validated by previous biochemical studies on SARS-CoV polymerase, which revealed their essential roles for stimulating the activity of nsp12 polymerase subunit (Subissi et al., 2014).

Based on the elongation complex of poliovirus polymerase (Gong and Peersen, 2010), we modeled the RNA template and product strands into the catalytic chamber of SARS-CoV-2 nsp12 subunit. This pseudo-elongation intermediate structure reveals the template entrance is supported by the finger extension loops and the finger-tip to guide the 3’-viral RNA (3’-vRNA) achieving the catalytic chamber (Figure 3D). The nucleotide-triphosphate (NTP) substrate enters through a channel at the back side of palm subdomain to reach the active site. The template and product stands form a duplex to exit the polymerase chamber in the front (Figure 3D). Since the viral genome and sub-genomic mRNA products are both functional in single-stranded form, it requires further steps assisted by other nsp subunits to separate the duplex for complete transcription and replication processes.

### The reduced activity of SARS-CoV-2 core polymerase complex

Given the residue substitutions between SARS-CoV-2 and SARS-CoV polymerase subunits albeit the high degree overall sequence similarity, we compared the enzymatic behaviors of the viral polymerases aiming to analyze their properties in terms of viral replication. Both sets of core polymerase complex could well mediate primer-dependent RNA elongation reactions templated by the 3’-vRNA. Intriguingly, the SARS-CoV-2 nsp12-nsp7-nsp8 complex displayed a much lower efficiency (~35%) for RNA synthesis as compared to the SARS-CoV counterpart (Figure 4A). As all three nsp subunits harbor some residue substitutions between the two viruses, we further conducted cross-combination analysis to evaluate the effects of each subunit on the efficiencies of RNA production. In the context of SARS-CoV-2 nsp12 polymerase subunit, replacement of the nsp7 cofactor subunit with that of SARS-CoV did not result in obvious effect on polymerase activity, whereas the introduction of SARS-CoV nsp8 subunit greatly boosted the activity to ~2.1 times of the homologous combination. Simultaneous replacement of the nsp7 and nsp8 cofactors further enhanced the efficiency for RNA synthesis to ~2.2 times of that for the SARS-CoV-2 homologous complex (Figure 4B). Consistent with this observation, the combination of SARS-CoV-2 nsp7-nsp8 subunits with the SARS-CoV nsp12 polymerase subunit compromised its activity as compared to the native cognate cofactors, among which the nsp8 subunit exhibited a more obvious effect than that for nsp7 (Figure 4C). These evidences suggested that the variations in nsp8 subunit rendered a significantly negative impact on the polymerase activity of SARS-CoV-2 nsp12. The non-significant effect of nsp7 on polymerase activity was quite conceivable as only one residue substitution occurred between the two viruses (Figure 2B). In addition, we also compared the polymerase activity of different nsp12 subunits in the same context of nsp7-nsp8 cofactors. Combined with either cofactor sets, the SARS-CoV-2 nsp12 polymerase showed a lower efficiency (~50%) for RNA synthesis as compared to the SARS-CoV counterpart (Figure 4D). This observation demonstrated that the residue substitutions in nsp12 also contributed to the reduction of its polymerase activity, with similar impact to the variations in the nsp8 cofactor.

**Figure 4.**
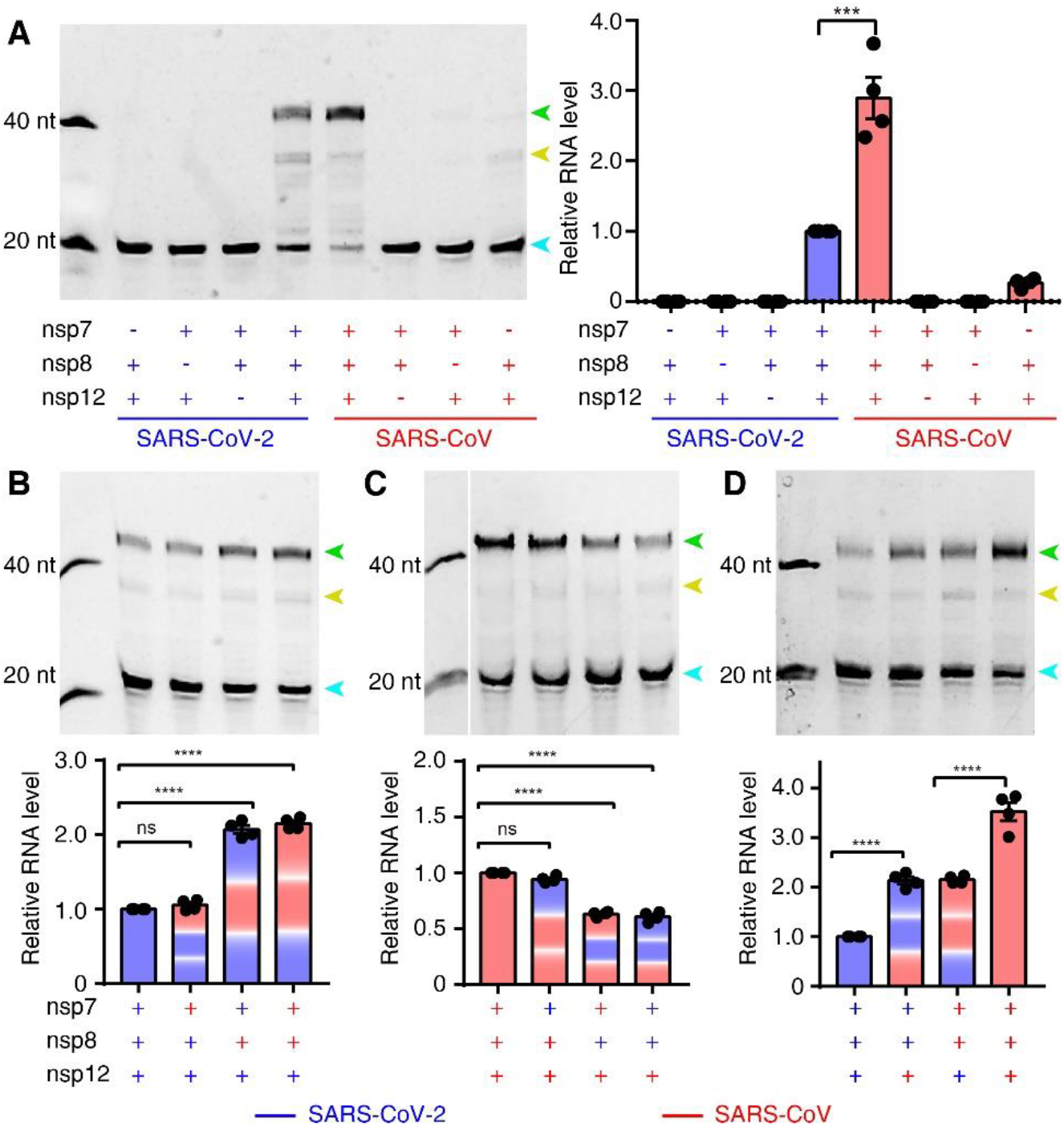
*In vitro* polymerase activity of nsp12 and regulatory effects of cofactors. **(A)** Comparison of RNA synthesis activities of SARS-CoV-2 and SARS-CoV core polymerase complex. The efficient activity of nsp12 polymerase requires the presence of both nsp7 and nsp8 cofactors. Apart from the fully elongated product (green arrowhead), some aberrant termination products were also observed (yellow arrowhead). The excess primer band is indicated by a cyan arrowhead. **(B, C)** Comparison of the regulatory effects of nsp7 and nsp8 cofactors in the context of SARS-CoV-2 **(B)** and SARS-CoV **(C)** nsp12 polymerase, respectively. **(D)** Comparison of the activity of nsp12 polymerase of different viruses in the same context of cofactors. The polymerase activity was quantified by integrating the intensity of the fully elongated product bands and the significance of difference was tested by one-way ANOVA based on the results of four independent experiments (n=4) using different protein preparations. *, *P*<0.05; **, *P*<0.01; ***, *P*<0.001; ****, *P*<0.0001. **See also Figure S4**.

### Impacts of amino acid substitutions on the core polymerase subunits

Despite that there are amino acid substitutions in all three subunits of the core polymerase complex between SARS-CoV-2 and SARS-CoV, none of these residues is located at the polymerase active site or the contacting interfaces between adjacent subunits (Figure 2B), suggesting these substitutions do not affect the inter-subunit interactions for assembly of the polymerase complex. To test this hypothesis, we measured the binding kinetics between different subunits of the two viruses by surface plasmon resonance (SPR) assays. Each interaction pair exhibited similar kinetic features for the two viruses, all with sub-micromolar range affinities (Figure 5A and B). We also tested the cross-binding between subunits of the two viruses, which revealed similar affinities for heterologous pairs as compared to the native homologous interactions (Figure 5C and D).

**Figure 5.**
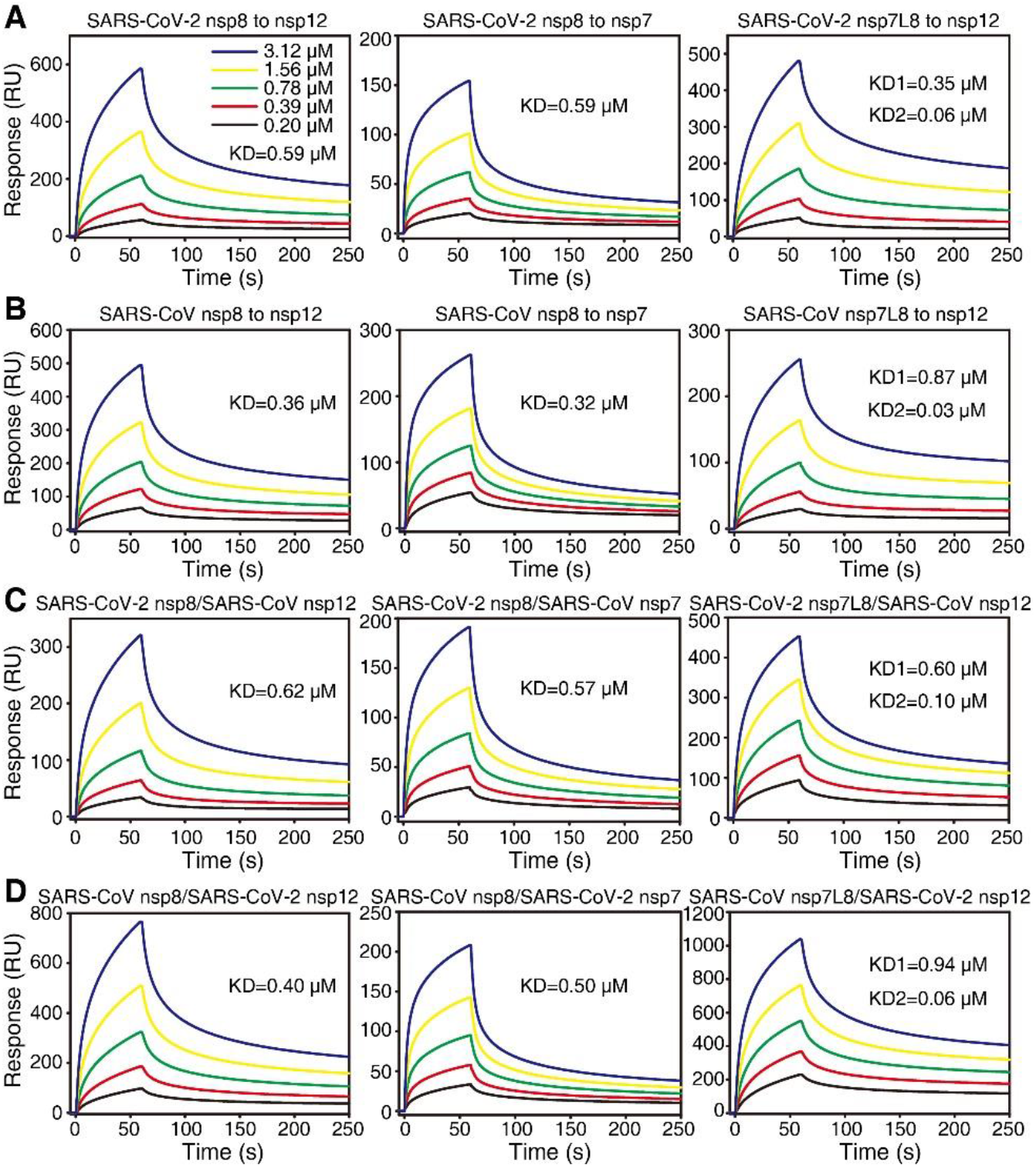
SPR binding kinetics of protein pairs in the polymerase complex of SARS-CoV-2 and SARS-CoV. **(A, B)** The binding profiles of homologous protein subunits of SARS-CoV-2 **(A)** and SARS-CoV **(B)**, respectively. **(C, D)** The cross-binding kinetics between protein subunits of the two viruses. All analytes were measured with serial-diluted concentrations as shown in **(A)**. The title is presented as analyte/immobilized ligand to facilitate comparison. The binding between nsp12 and nsp7L8 fusion protein was fitted with the heterogeneous binding mode as the nsp7-nsp8 heterodimer exhibited non-uniform conformations in solution (Zhai et al., 2005). It can occupy either cofactor binding site as stable nsp7-nsp8 complex or free nsp8 once the nsp7 detach from the heterodimer. Both equilibrium binding constant values (KD1 and KD2) were calculated in this mode. The data shown is a representative result of three independent experiments using different protein preparations, all of which produced similar results. **See also Figure S4**.

We then compared the thermostability of each component in the polymerase complex of the two viruses (Figure 6). Consistent with the almost identical sequences, the nsp7 of both viruses displayed comparable melting behaviors in the circular dichroism (CD) profiles, demonstrating similar thermostabilities of the two proteins (Figure 6A and D). In contrast, both the nsp8 and nsp12 subunits of SARS-CoV-2 showed lower melting temperature (Tm) values as compared to the corresponding subunits of SARS-CoV, suggesting the poorer thermostability of SARS-CoV-2 proteins (Figure 6B, C, E and F).

**Figure 6.**
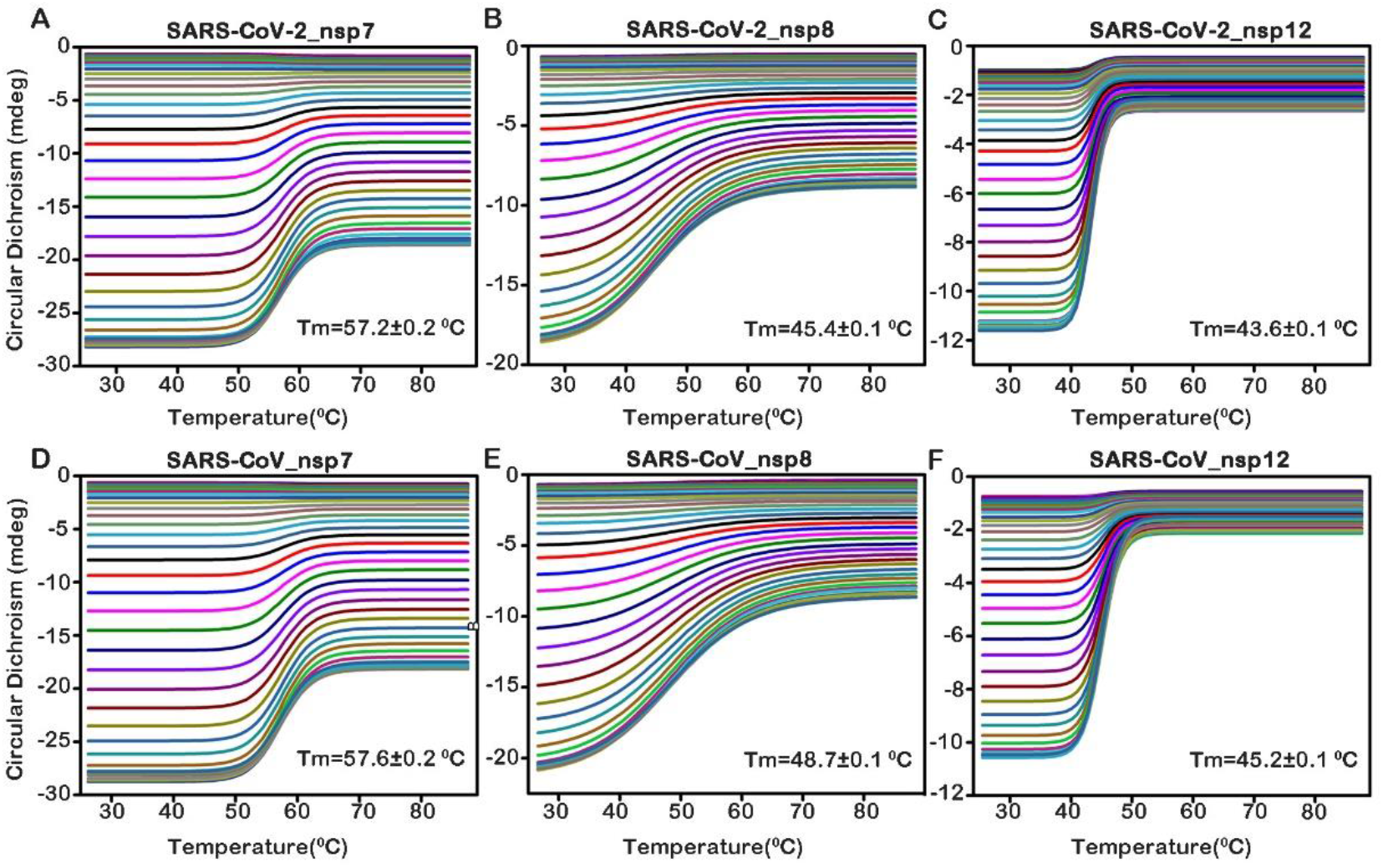
CD profiles of individual subunits in the core polymerase complex. The multi-wavelength (215-260 nm) CD spectra of SARS-CoV-2 **(A-C)** and SARS-CoV **(D-F)** polymerase components at different temperatures. The Tm values were calculated to evaluate the thermostability of each protein subunit. The data shown are representative results of two independent experiments using different protein preparations. **See also Figure S4**.

Taken together, the residue substitutions in SARS-CoV-2 nsp12 polymerase subunit and nsp7-nsp8 cofactors compromise the efficiency of RNA synthesis by the core polymerase complex and reduce the thermostability of individual protein subunits as compared to the counterparts of SARS-CoV. These changes may indicate the adaptive evolution of SARS-CoV-2 towards the human hosts with a relatively lower body temperature than bats which are potentially the natural host of a panel of zoonotic viruses, including both SARS-CoV and SARS-CoV-2 (O’Shea et al., 2014; Zhou et al., 2020).

## Discussion

The structural information of coronavirus polymerase interaction with cofactors suggests a common theme of viral RdRp activation despite being executed by different structural components. The coronavirus polymerase subunit requires multiple cofactors/subunits for complete transcription and replication functions, similar to the related flaviviruses which also harbor a positive-sense RNA genome (Aktepe and Mackenzie, 2018; Sevajol et al., 2014; Ziebuhr, 2005). In contrast, the sNSVs utilize fewer multi-subunit protein components to accomplish similar processes, which could be activated by RNA segments instead of proteins (Gerlach et al., 2015; Peng et al., 2020; Pflug et al., 2014). As revealed by the coronavirus core polymerase structures, it lacks the essential component for unwinding the template-product hybrid which is required to release the single-strand mRNA and viral genome for protein expression and virion assembly. In the structure of sNSV polymerases, a lid domain is present at the intersection region of template and product exit tunnels to force duplex deformation before leaving the polymerase chamber (Gerlach et al., 2015; Peng et al., 2020; Reich et al., 2014). The nsp13 subunit has been shown with RNA helicase activity, suggesting its involvement in RNA synthesis at the post-catalytic stage (Adedeji et al., 2012). Further investigations are required to understand how this process takes place.

Of note, we demonstrate the amino acid substitutions in the polymerase and cofactors of SARS-CoV-2 lead to obviously reduced activity for RNA synthesis as compared to SARS-CoV core polymerase complex. Indeed, these observations are based on partial components of the multiple-subunit holoenzyme for coronavirus replication which also involves proofreading and capping by other nsp subunits, e.g. the nsp10-nsp14 exonuclease subcomplex, nsp13 RNA 5’-triphosphatase, and the nsp14 and nsp16 methyltransferases (Sevajol et al., 2014; Ziebuhr, 2005). These steps would also render important determinants for the efficiency and accuracy of RNA synthesis by coronavirus replication machinery. Thus, the collective behavior of SARS-CoV-2 polymerase complex in the context of an authentic viral replication cycle still remains an open question to be further explored. On the other hand, the lower thermostability of SARS-CoV-2 polymerase subunits indicate its well adaptation for humans which have a relatively lower body temperature compared to bats, the potential natural host of SARS-CoV-2 (O’Shea et al., 2014; Zhou et al., 2020). Interestingly, we also found that the closely-related bat coronavirus RaTG13 showed an extremely high sequence identity of core polymerase subunits to SARS-CoV-2, in which the nsp7 and nsp8 cofactors are strictly identical and the nsp12 catalytic subunit harbors only four residue replacements between the two viruses (Figure S5), suggesting similar enzymatic properties and thermostabilities of their polymerase components. This observation indicated the RaTG13 coronavirus has been well adapted to human hosts in terms of viral replication machinery and might further support the probable bat-origin of SARS-CoV-2 (Zhou et al., 2020).

In summary, our structural and biochemical analyses on SARS-CoV-2 core polymerase complex improve our understanding on the mechanisms of RNA synthesis by different viral RdRps and highlight a common theme for polymerase activation by stabilizing critical catalytic motifs via diverse means. In addition, the different biochemical properties of polymerase components of SARS-CoV-2 and SARS-CoV suggest the clues for adaptive evolution of coronaviruses in favor of human hosts.

## Acknowledgements

We thank all staff members in the Center of Biological Imaging (CBI), Institute of Biophysics (IBP), Chinese Academy of Sciences (CAS), for assistance with data collection. We are grateful to the Core Facility in Institute of Microbiology, Chinese Academy of Sciences (CAS) for assistance in SPR experiments. This study was supported by the Strategic Priority Research Program of CAS (XDB29010000), the National Science and Technology Major Project (2018ZX10101004), National Key Research and Development Program of China (2020YFC0845900), the National Natural Science Foundation of China (NSFC) (82041016, 81871658 and 81802010), and a grant from the Bill & Melinda Gates Foundation. M.W. is supported by the National Science and Technology Major Project (2018ZX09711003) and National Natural Science Foundation of China (NSFC) (81802007). R.P. is supported by the Young Elite Scientist Sponsorship Program (YESS) by China Association for Science and Technology (CAST) (2018QNRC001). Y.S. is also supported by the Excellent Young Scientist Program and from the NSFC (81622031) and the Youth Innovation Promotion Association of CAS (2015078).

## Author contributions

Y.S. conceived the study. Q.P. J.Z., B.Y., M.W., X.W., Y.Sun and Q.W. purified the protein samples and conducted biochemical studies. Q.P. and R.P. performed cryo-EM analysis. R.P. and J.Q. built the atomic model. Q.P., R.P., M.W. and Y.S. analyzed the data and wrote the manuscript. All authors participated in the discussion and manuscript editing. Q.P., R.P., B.Y. and J.Z. contributed equally to this work.

## STAR Methods

### Plasmids, bacterial strains and cell lines

The nsp12 polymerase subunit of both SARS-CoV and SARS-CoV-2 were cloned into the pFastbac-1 vector and expressed in the High Five insect cell line. The nsp7, nsp8 and nsp7L8 fusion protein were incorporated into the pET-21a plasmid and expressed in *E. Coli* BL21 (DE3) strain.

### Protein expression and purification

The codon-optimized sequences of nsp7 and nsp8 were synthesized with N-terminal 6×histidine tag and inserted into pET-21a vector for expression in *E. coli* (Synbio Tec, Suzhou, China). For the nsp7L8 fusion protein, the sequence was also codon-optimized for *E. Coli* expression system and a 6×histidine linker was introduced between the nsp7 and nsp8 subunits (Genewiz Tec, Suzhou, China). Protein production was induced with 1 mM isopropylthio-galactoside (IPTG) and incubated for 14-16 hours at 16 °C. Bacterial cells were harvested by centrifugations (12,000 rpm, 10 min), resuspended in buffer A (20 mM HEPES, 500 mM NaCl, 2 mM Tris (2-carboxyethyl) phosphine (TCEP), pH 7.5, and lysed by sonication. Cell debris were removed via centrifugation (12,000 rpm, 1h) and filtration with a 0.22 μm cut-off filter. The supernatant was loaded onto a HisTrap column (GE Healthcare) for initial affinity purification. The target proteins were eluted using buffer A supplemented with 300 mM imidazole. Fractions were pooled and subjected to size-exclusion chromatography (SEC) with a Superdex 200 increase column (GE Healthcare). The final product was concentrated and stored at −80 °C.

For nsp12 proteins, the genes were codon-optimized for *Spodoptera frugiperda* incorporated into the pFastBac-1 plasmid with a C-terminal thrombin proteolysis site, a 6× histidine and two tandem Strep tags. Proteins were expressed with High Five cells at 27 °C for 48 h post infection. Cells were collected by centrifugation (3,000 rpm, 10 min) and resuspended in buffer B (25 mM HEPES, 300 mM NaCl, 1 mM MgCl_2_, and 2mM TCEP, pH 7.4. The cell suspension was lysed by sonication and the lysate was clarified using ultracentrifugation (30,000 rpm, 2 h) and filtered with 0.22 μm cut-off members. The resulting supernatant was applied to a StrepTrap column (GE Healthcare) to capture the target proteins. The bound proteins were eluted with buffer B supplemented with 2.5 mM desthiobiotin. Target fractions were pooled and subjected to further purification by SEC using a Superdex 200 increase column (GE Healthcare). The final product was concentrated and stored at −80 °C before use.

### Cryo-EM sample preparation and data collection

An aliquot of 3 μL protein solution (0.6 mg/mL) was applied to a glow-discharged Quantifiol 1.2/1.3 holey carbon grid and blotted for 2.5 s in a humidity of 100% before plunge-freezing with an FEI Vitrobot Mark IV. Cryo-samples were screened using an FEI Tecnai TF20 electron microscope and transferred to an FEI Talos Arctica operated at 200 kV for data collection. The microscope was equipped with a post-column Bioquantum energy filter (Gatan) which was used with a slit width of 20 eV. The data was automatically collected using SerialEM software (http://bio3d.colorado.edu/SerialEM/). Images were recorded with a Gatan K2-summit camera in super-resolution counting mode with a calibrated pixel size of 0.8 Å at the specimen level. Each exposure was performed with a dose rate of 10 e^−^/pixel/s (approximately 15.6 e^−^/Å^2^/s) and lasted for 3.9 s, resulting in an accumulative dose of ~60 e^−^/Å^2^ which was fractionated into 30 movie-frames. The final defocus range of the dataset was approximately −1.4 to −3.4 μm.

### Image processing

The image drift and anisotropic magnification was corrected using MotionCor2 (Zheng et al., 2017). Initial contrast transfer function (CTF) values were estimated with CTFFIND4.1 (Rohou and Grigorieff, 2015) at the micrograph level. Images with an estimated resolution limit worse than 5 Å were discarded. Particles were automatically picked with RELION-3.0 (Zivanov et al., 2018) following the standard protocol. In total, approximately 1,860,000 particles were picked from ~4,200 micrographs. After 3 rounds of extensive 2D classification, ~924,000 particles were selected for 3D classification with the density map of SARS-CoV nsp12-nsp7-nsp8 complex (EMDB-0520) as the reference which was low-pass filtered to 60 Å resolution. After two rounds of 3D classification, a clean subset of ~101,000 particles was identified, which displayed clear features of secondary structural elements. These particles were subjected to 3D refinement supplemented with per-particle CTF refinement and dose-weighting, which led to a reconstruction of 3.65 Å resolution estimated by the gold-standard Fourier shell correlation (FSC) 0.143 cut-off value. The local resolution distribution of the final density map was calculated with ResMap (Kucukelbir et al., 2014).

### Model building and refinement

The structure of SARS-CoV nsp12-nsp7-nsp8 complex (PDB ID: 6NUR) was rigidly docked into the density map using CHIMERA (Pettersen et al., 2004). The model was manually corrected for local fit in COOT (Emsley et al., 2010) and the sequence register was corrected based on alignment. The initial model was refined in real space using PHENIX (Adams et al., 2010) with the secondary structural restraints and Ramachandran restrains applied. The model was further adjusted and refined iteratively for several rounds aided by the stereochemical quality assessment using MolProbity (Chen et al., 2010). The representative density and atomic models are shown in Figure. S3. The statistics for image processing and model refinement are summarized in the Table S1. Structural figures were rendered by either CHIMERA (Pettersen et al., 2004) or PyMOL (https://pymol.org/).

### *In vitro* polymerase activity assay

The activity of SARS-CoV-2 polymerase complex was tested as previously described for SARS-CoV nsp12 with slight modifications. Briefly, a 40-nt template RNA (5’-CUAUCCCCAUGUGAUUUUAAUAGCUUCUUAGGAGAAUGAC-3’, Takara) corresponding to the 3’end of the SARS-CoV-2 genome was annealed to a complementary 20-nt primer containing a 5’-fluorescein label (5’FAM-GUCAUUCUCCUAAGAAGCUA-3’, Takara). To perform the primer extension assay, 1 μM nsp12, nsp7 and nsp8 were incubated for 30 min at 30 °C with 1 μM annealed RNA and 0.5 mM NTP in a reaction buffer containing 10 mM Tris-HCl (pH 8.0), 10 mM KCl, 1 mM beta-mercaptoethanol and 2 mM MgCl2 (freshly added prior usage). The products were denatured by boiling (100 °C, 10 min) in the presence of formamide and separated by 20% PAGE containing 9 M urea run with 0.5×TBE buffer. Images were taken using a Vilber Fusion system and analyzed with the Image J software.

### SPR assay

The affinities between nsp12, nsp7 and nsp8 or nsp7L8 proteins were measured at room temperature (r.t.) using a Biacore 8K system with CM5 chips (GE Healthcare). The nsp12 protein was immobilized on the chip with a concentration of 100 μg/mL (diluted by 0.1 mM NaAc, pH 4.0), and the nsp7 protein was immobilized with a concentration of 50 μg/mL (diluted by 0.1 mM NaAc, pH 4.5). For all measurements, the same running buffer was used which consists of 20 mM HEPES, pH 7.5,150 mM NaCl and 0.005% tween-20. Proteins were pre-exchanged into the running buffer by SEC prior to loading to the system. A blank channel of the chip was used as the negative control. Serially diluted protein solutions were then flowed through the chip surface. The Multi-cycle binding kinetics was analyzed with the Biacore 8K Evaluation Software (version1.1.1.7442) and fitted with a two-state reaction binding model (for ligand nsp8) or heterogeneous ligand binding model (for ligand nsp7L8).

### Circular dichroism (CD) measurement

The thermostability of nsp12, nsp7 and nsp8 were tested by measuring the CD spectra of each protein at different temperatures. The multi-wavelength (215-260 nm) CD spectra of each protein were recorded with a Chirascan spectrometer (Applied Photophysics) using a thermostatically controlled cuvette at rising temperatures from 25 to 99 °C with 0.5 °C intervals and an elevating rate of 1 °C/min. The data was analyzed using Global3 software and the Tm values were calculated for each sample.

### Data availability

The cryo-EM density map and atomic coordinates have been deposited to the Electron Microscopy Data Bank (EMDB) and the Protein Data Bank (PDB) with the accession codes EMD-30226 and 7BW4, respectively. All other data are available from the authors on reasonable request.

## Supplemental Information

**Figure S1.**
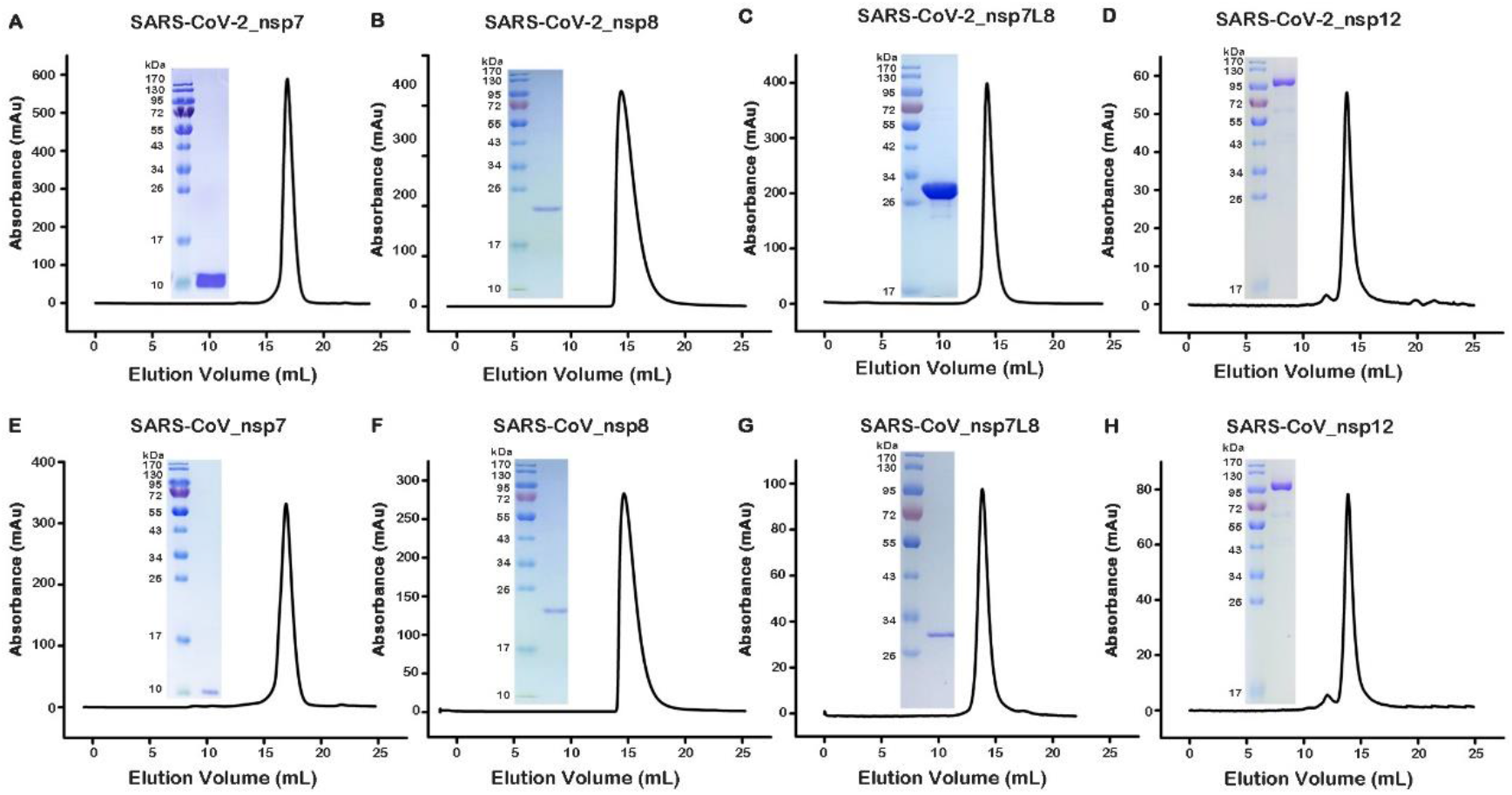
Purification of nsp12 polymerase and nsp7-nsp8 cofactors. Size-exclusion chromatography and SDS-PAGE profiles of the nsp7 **(A, E)**, nsp8 **(B, F)**, nsp7L8 (ns7-nsp8 heterodimer) **(C, G)** and nsp12 **(D, H)** proteins of SARS-CoV-2 and SARS-CoV. All these protein samples showed good behaviors in solution with high homogeneity and purity. **Related to Figure 1**.

**Figure S2.**
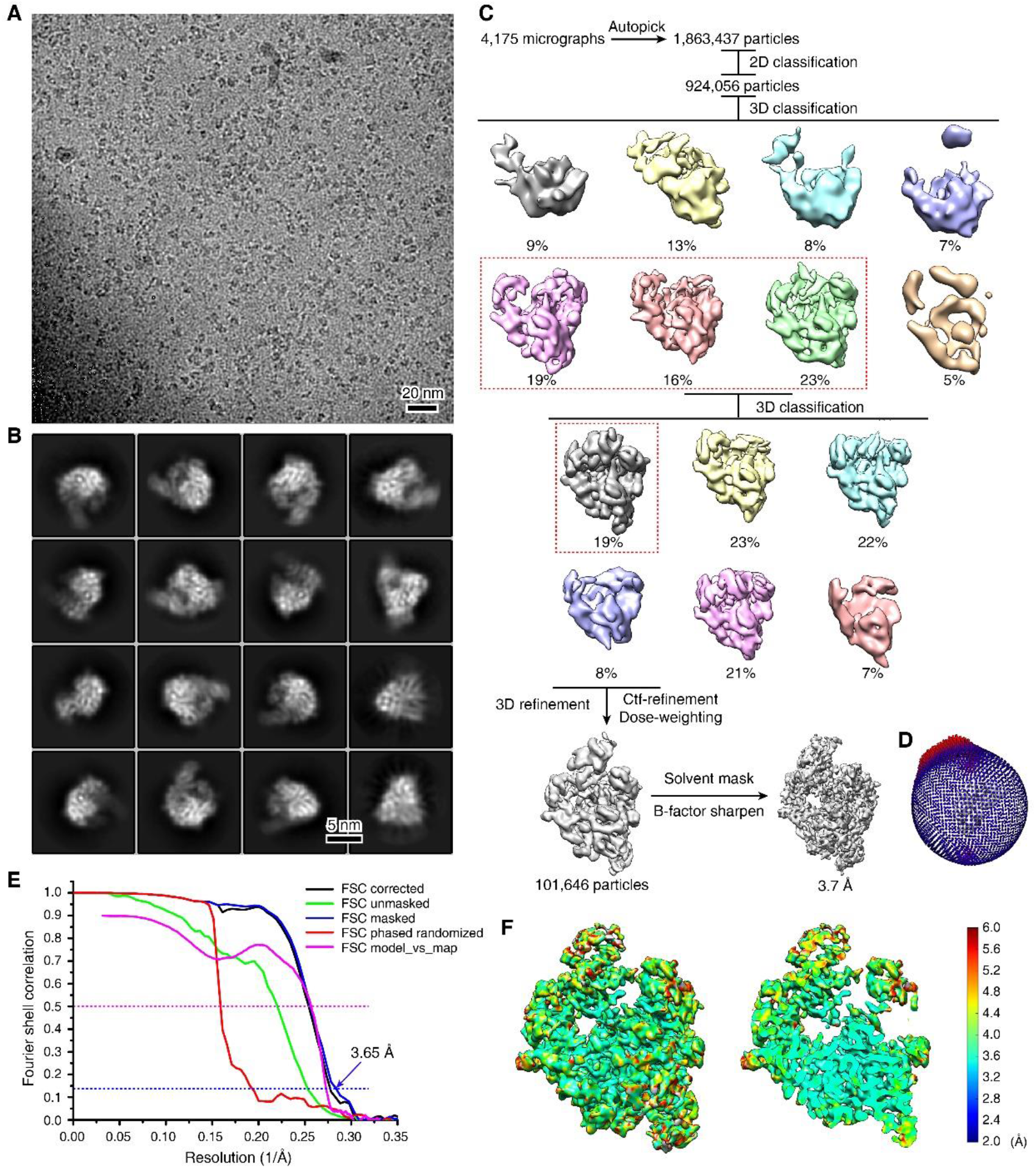
Cryo-EM analysis of SARS-CoV-2 core polymerase complex. **(A)** A representative micrograph of SARS-CoV-2 nsp12-nsp7-nsp8 complex (out of >4,000 micrographs). **(B)** Typical 2D class average images. Three rounds of 2D classification were performed. **(C)** Image processing workflow. The selected classes for next-step processing are indicated by red dashed boxes. **(D)** Euler angle distribution of the final reconstruction. **(E)** FSC curves for evaluating the resolution of final reconstruction and model-map correlations. **(F)** Local resolution assessment of the final density map. **Related to Figure 1**.

**Figure S3.**
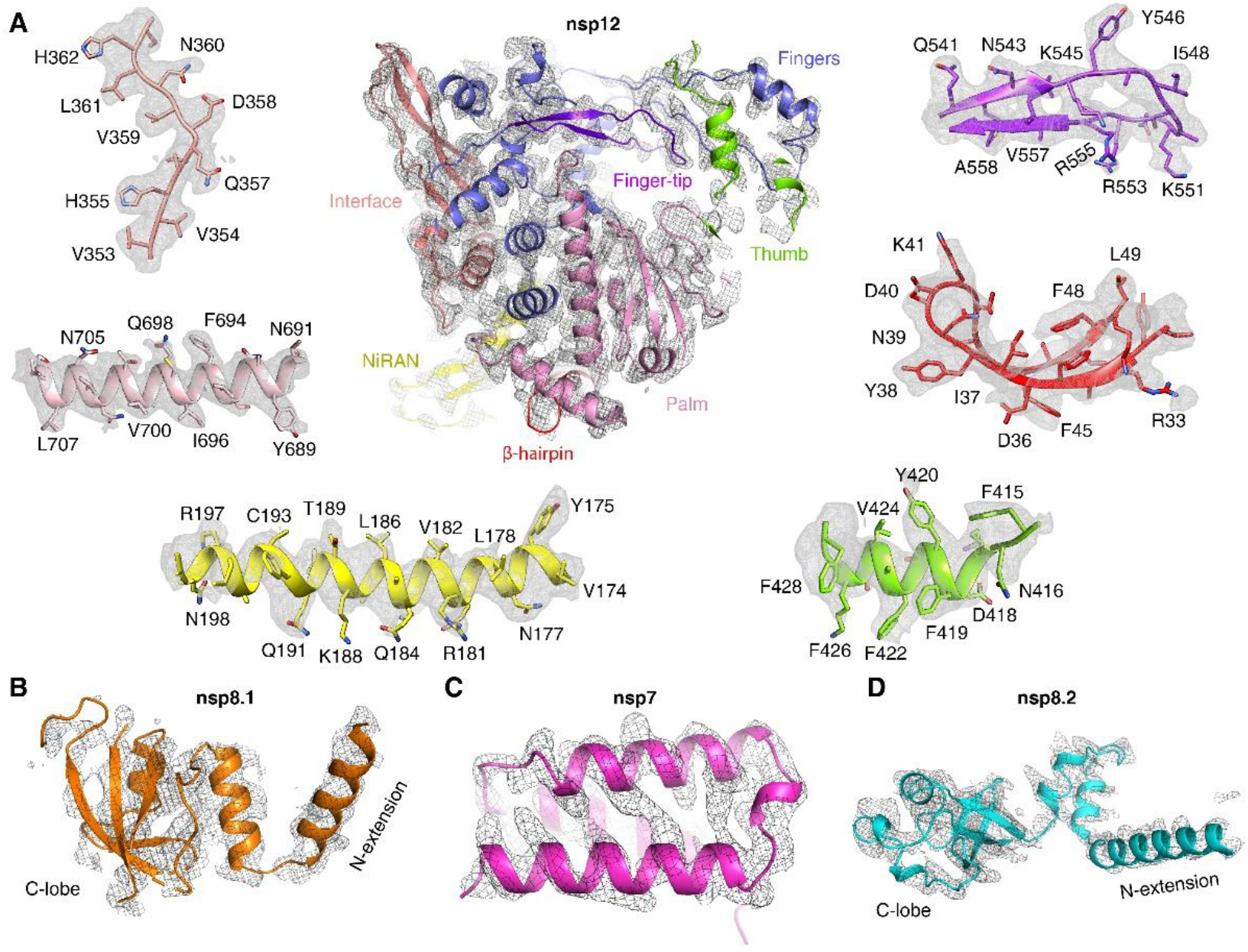
Representative density and atomic models of individual polymerase components. **(A)** Overall density and atomic model of the nsp12 subunit and the close-up views in selected regions. Most bulky side chains could be clearly resolved in the density map. **(B-D)** The density maps for nsp7 and nsp8 individual cofactor subunits. **Related to Figure 1**.

**Figure S4.**
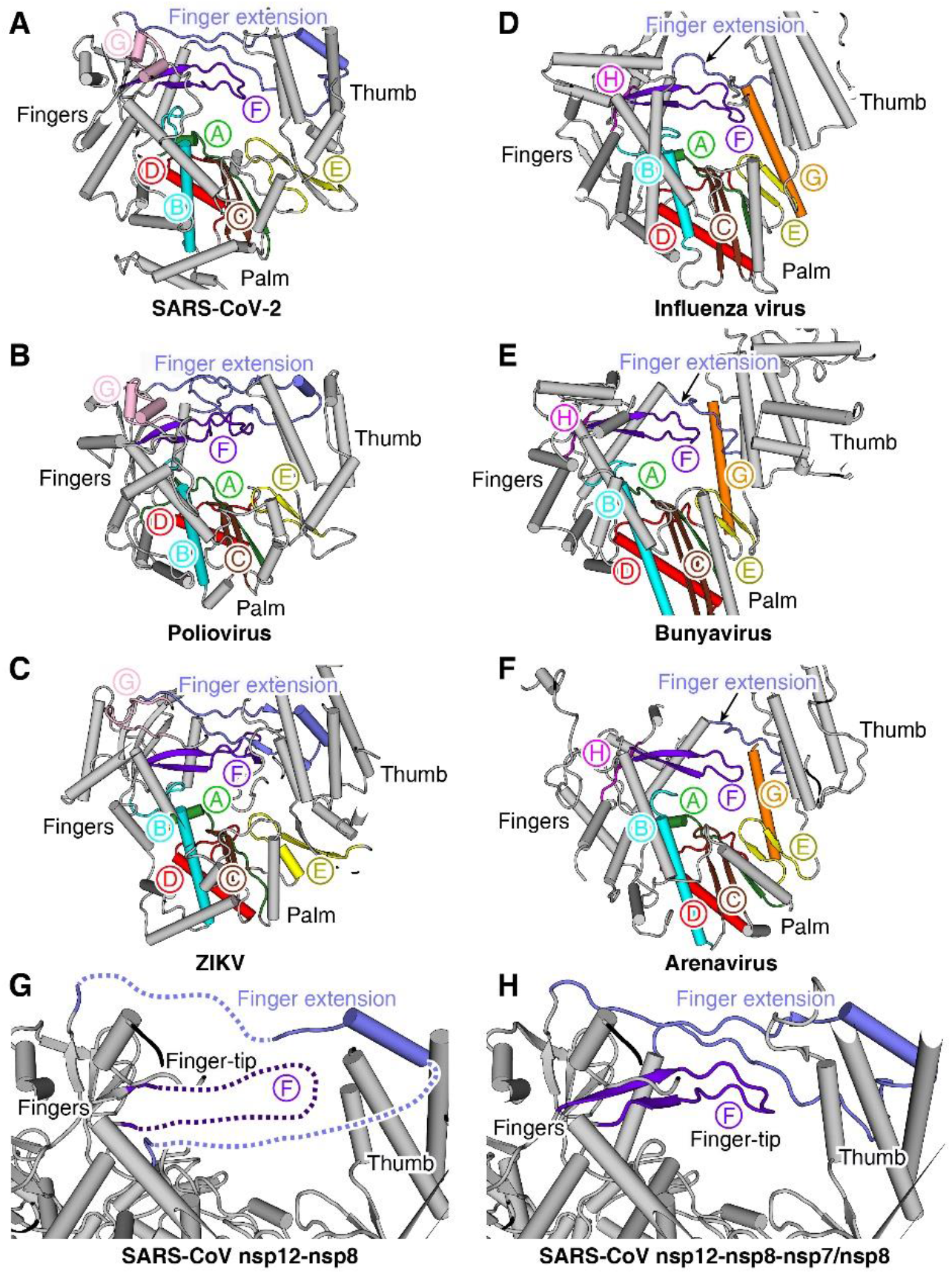
Structural comparison of different viral RdRps. (A-F) Comparison of catalytic motifs in different viral RdRps. The SARS-CoV-2, poliovirus (PDB ID: 3OL8) and ZIKV (PDB ID: 5WZ3) belong to positive-sense RNA virus group. The influenza virus (PDB ID: 4WRT), bunyavirus (exemplified by the La Crosse orthobunyavirus, LACV; PDB ID: 5AMQ) and arenavirus (exemplified by the Lassa virus, LASV; PDB ID: 6KLC) **(F)** belong to sNSV group. Each catalytic motif is represented by a unique color. The finger extension is highlighted with a black arrow in each structure. **(G, H)** The structures of SARS-CoV nsp12 polymerase with **(H)** or without **(G)** nsp7-nsp8 heterodimer binding. The finger-tip (motif F) and finger extension loops is disordered on its own, which are significantly stabilized by nsp7-nsp8 cofactors binding. **Related to Figures 2-6**.

**Figure S5.**
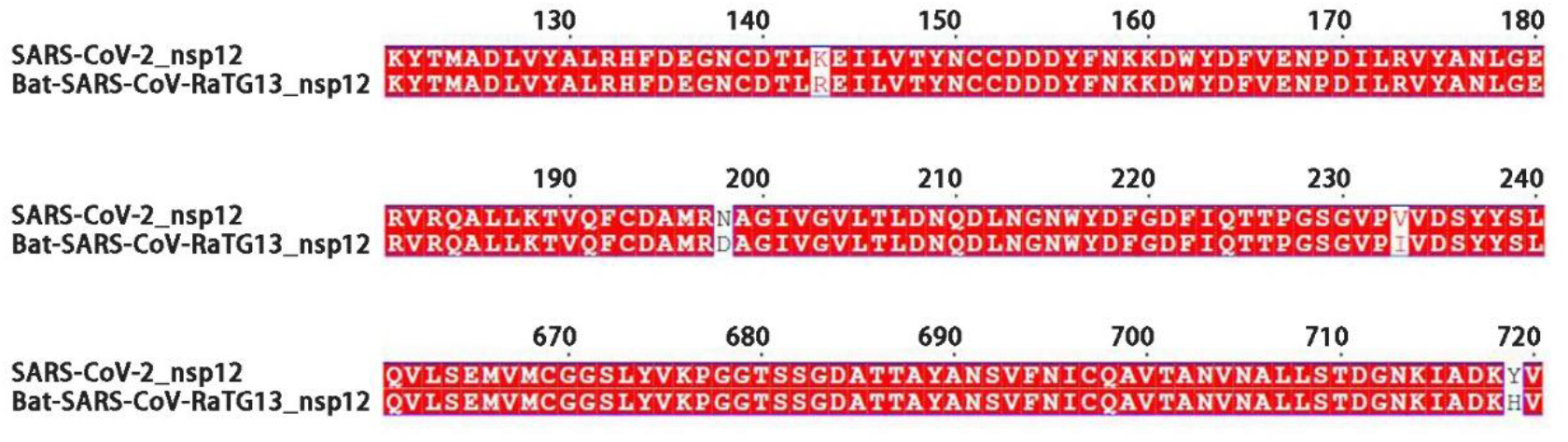
Sequence alignment of nsp12 polymerase subunit in selected regions of SARS-CoV-2 and RaTG13 bat coronavirus. The four residue substitutions between the two proteins are highlighted by white background. **Related to Figure 6**.

**Table S1.** Cryo-EM data processing and refinement statistics.

| SARS-CoV-2 nsp12-nsp7-nsp8 complex |  |
| --- | --- |
| **Data collection and processing** |  |
| Magnification | 62,500 |
| Voltage (kV) | 200 |
| Electron exposure (e<sup>-</sup>/Å<sup>2</sup>) | 60 |
| Defocus range (μm) | -1.4 to -3.4 |
| Pixel size (Å) | 0.8 |
| Symmetry imposed | C1 |
| Final particle images (no.) | 101,646 |
| Map resolution (Å) | 3.65 |
| FSC threshold | 0.143 |
| Map resolution range (Å) | 3.0-6.0 |
| **Refinement** |  |
| Initial model used (PDB code) | 6NUR |
| Model resolution range (Å) | Up to 3.7 |
| Map sharpening *B* factor (Å<sup>2</sup>) | -135 |
| Map correlation coefficient |  |
| Whole unit cell | 0.77 |
| Around atoms | 0.77 |
| Model composition |  |
| Non-hydrogen atoms | 8,595 |
| Protein residues | 1,078 |
| *B* factors (Å<sup>2</sup>) |  |
| Protein | 89 |
| Ligand | 74 |
| R.m.s. deviations |  |
| Bond lengths (Å) | 0.004 |
| Bond angles (°) | 0.65 |
| **Validation** |  |
| MolProbity score | 2.21 |
| Clashscore | 15.73 |
| Poor rotamers (%) | 1.68 |
| Ramachandran plot |  |
| Favored (%) | 95.28 |
| Allowed (%) | 4.53 |
| Disallowed (%) | 0.19 |

